# Can biomass distribution across trophic levels predict trophic cascades?

**DOI:** 10.1101/2020.04.04.025460

**Authors:** Núria Galiana, Jean-François Arnoldi, Matthieu Barbier, Amandine Acloque, Claire de Mazancourt, Michel Loreau

## Abstract

The biomass distribution across trophic levels (biomass pyramid), and cascading responses to perturbations (trophic cascades), are archetypal representatives of the interconnected set of static and dynamical properties of food chains. A vast literature has explored their respective ecological drivers, sometimes generating correlations between them. Here we instead reveal a fundamental connection: both pyramids and cascades reflect the dynamical sensitivity of the food chain to changes in species intrinsic rates. We deduce a direct relationship between cascades and pyramids, modulated by what we call trophic dissipation – a synthetic concept that encodes the contribution of top-down propagation of consumer losses in the biomass pyramid. Predictable across-ecosystem patterns emerge when systems are in similar regimes of trophic dissipation. Data from 31 aquatic mesocosm experiments demonstrate how our approach can reveal the causal mechanisms linking trophic cascades and biomass distributions, thus providing a road map to deduce reliable predictions from empirical patterns.

## 1 Introduction

Food chains are a central concept of ecology, providing intuitions and predictions about basic static properties of food webs, such as the distribution of biomass across the trophic hierarchy, i.e. the trophic pyramid (Lindeman, 1942; Elton, 1927), but also about more complex dynamical processes, such as the propagation of a perturbation along the chain, i.e. trophic cascades (Shurin *et al*., 2002; Borer *et al*., 2005; Carpenter *et al*., 1985; Polis *et al*., 2000). Trophic cascades are prevalent in nature, yet of highly variable strength across systems (Polis *et al*., 2000; Shurin *et al*., 2002; Frank *et al*., 2006) and thus hard to predict, an observation that has prompted the proliferation of studies exploring the mechanisms behind their existence and magnitude (Halaj & Wise, 2001; Shurin & Seabloom, 2005; Borer *et al*., 2005; Heath *et al*., 2014; Ripple *et al*., 2016).

Studies of static and dynamical properties of food chains have historically developed in isolation, generating different lines of investigation, which have seldom been connected rigorously (but see Jonsson (2017); McCauley *et al*. (2018); Barbier & Loreau (2019); Rossberg *et al*. (2019)). A number of applied studies have proposed to predict an ecosystem’s “health” and dynamical response from its more accessible static biomass pyramid or size spectrum (Shin *et al*., 2005; Cury *et al*., 2005). Yet others claim that such static features are not informative about the underlying food chain dynamics (Trebilco *et al*., 2016; McCauley *et al*., 2018; Woodson *et al*., 2018). Here, we propose that there are several ways in which cascades and biomass pyramids could be related (or not), leading to contrasting interpretations. They can correlate across ecosystems via dependency on common parameters, as suggested by recent theory (Heath *et al*., 2014; Barbier & Loreau, 2019, Figure 1a). But, as we will argue in depth, they can also be more tightly connected, when the two phenomena arise from a single dynamical mechanism that acts as a common proximate cause (Figure 1b). Importantly, a direct link is required to reliably predict dynamical behaviors from static snapshots alone, without additional knowledge of potential confounding factors.

**Figure 1:**
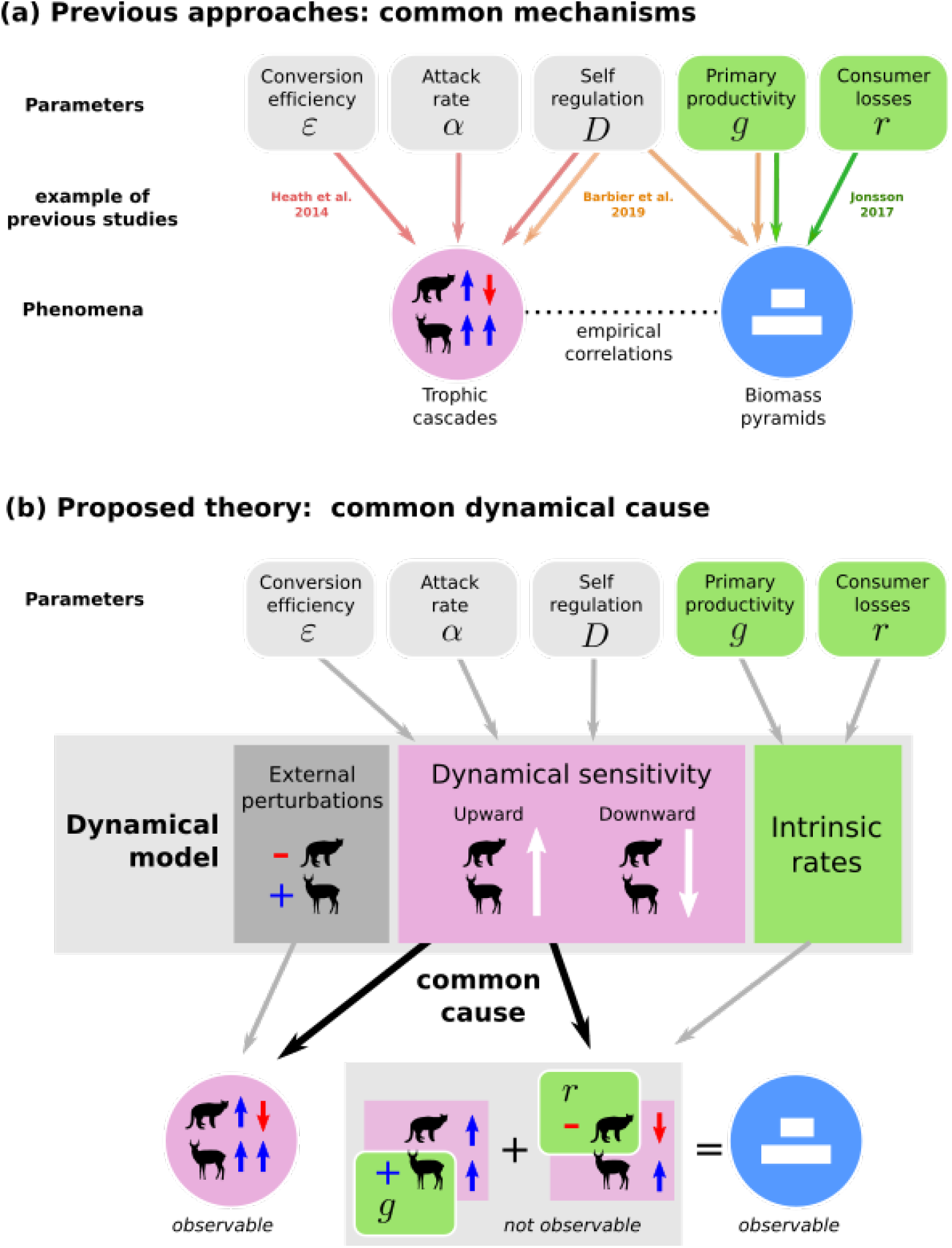
Relating biomass pyramids and trophic cascades. (a) Previous studies have investigated ecological parameters that affect the pyramidal distribution of biomass across trophic levels and the strength of trophic cascades, e.g. (Heath *et al.,* 2014; Jonsson, 2017; Barbier & Loreau, 2019) (arrows are not exhaustive). These parameters may drive empirical correlations between the two phenomena. (b) Here we focus on a direct common cause: the dynamical sensitivity of the ecosystem to both upward and downward effects along the food chain. This dynamical sensitivity combines with external perturbations to create trophic cascades, and with the food chain’s intrinsic rates of biomass gain and loss *(g* and *r)* to determine biomass pyramids. By factoring out perturbation intensity, trophic cascades readily provide an estimate of dynamical sensitivity. Biomass pyramids, however, are the result of a complex entanglement of upward and downward effects, which cannot be directly observed and separated. Across ecosystems, a one-to-one relationship between biomass pyramids and trophic cascades exists if the main source of variation is dynamical sensitivity (see Figure 2).

This dichotomy is at the core of the present study. We specifically ask when the shape of the standing biomass pyramid is -or is not-*causally* related to the strength of trophic cascades. In other words, we investigate the nature and drivers of *the relationship* between cascade strength and the shape of the biomass pyramid, to then determine when and how the latter can be used to predict the former. This novel perspective complements previous studies: we do not ask why the cascades are weak or strong (Borer *et al*., 2005), but rather why their strength can or cannot be predicted from the observation of the biomass pyramid.

The simplest expectation, for any ecosystem with multiple compartments, is that the response of a compartment to a perturbation is proportional to its standing biomass. We take it as our baseline (Figure 2a). This expectation may appear intuitive, yet many classical results on food chains do not, in fact, support it. On a quantitative level, deviations from strict proportionality occur in many theoretical and empirical studies, and have been used to define whether a cascade is strong or weak (Hedges *et al*., 1999; Shurin *et al*., 2002). These deviations are commonly interpreted as an amplification or attenuation of perturbations as they propagate along the chain (Box 1). The proportionality hypothesis implies that *both* upward and downward effects are directly related to the biomass distribution. Yet, it has long been noticed that the significance of these two types of effects can widely differ across systems. Each has been the focus of a classic perspective on food chain structure and dynamics. The resource-based perspective posits that producers determine the biomass of higher trophic levels (Elton, 1927; Lindeman, 1942), while the consumer-based perspective emphasizes the role of predators in regulating the biomass of lower trophic levels (Hairston *et al*., 1960; Leibold, 1996; Gruner *et al*., 2008).

**Figure 2:**
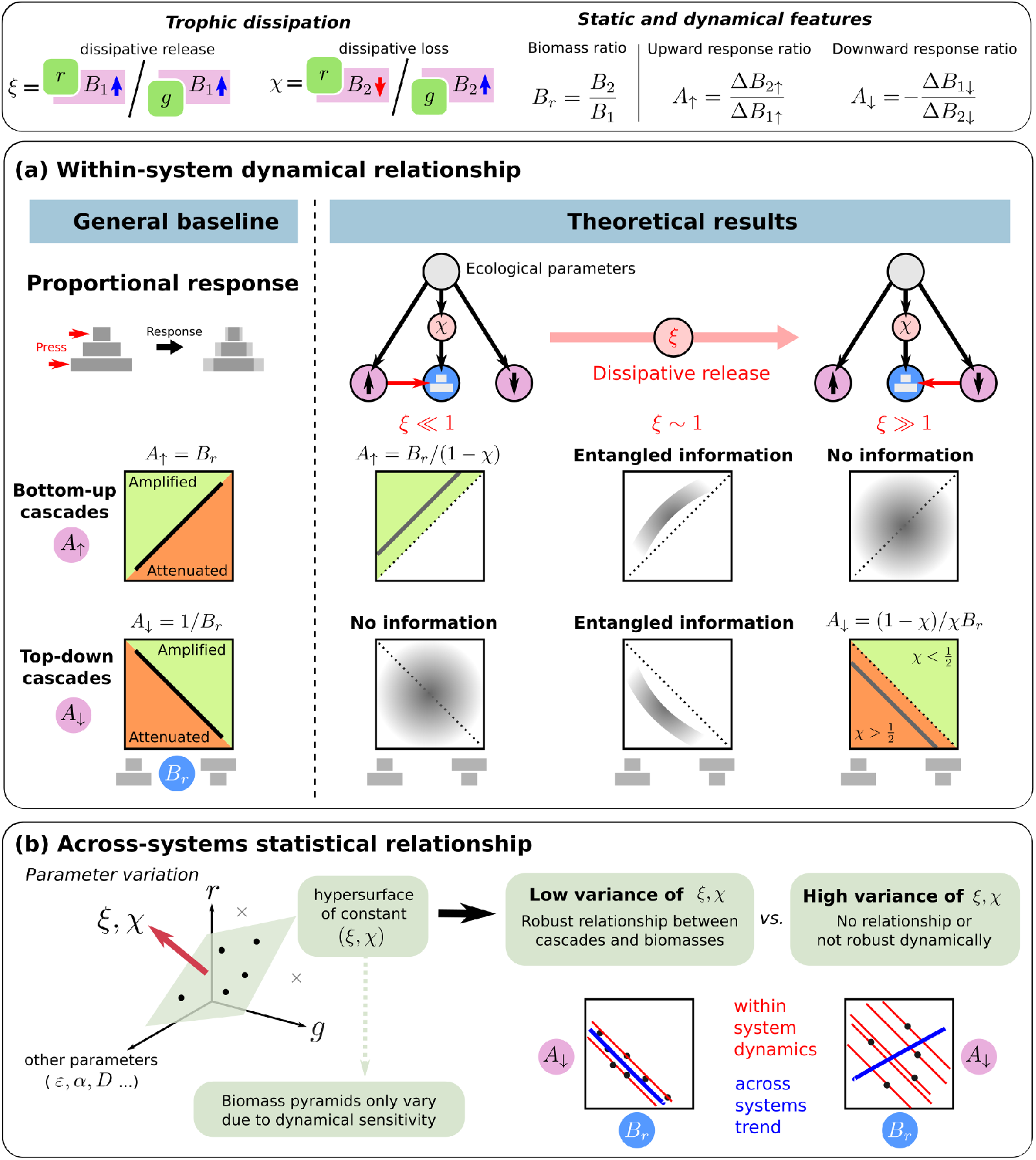
Relationship between static and dynamic properties in food chains. (a) Within one system, the simplest expectation is that a trophic level’s response to a perturbation is proportional to its biomass. Deviations from this trend indicate the amplification or attenuation of perturbations between levels (Shurin *et al*., 2002). Food chain theory instead suggests two limiting cases where equilibrium biomasses are related either to bottom-up effects only, or to top-down effects only. Our framework connects these two limits on a single axis of *trophic dissipation.* From the consumer perspective we identify *dissipative loss χ* and from the resource perspective *dissipative release ξ* (see Box 1). When these two parameters are fixed, all other sources of variation affect biomasses only through the same dynamical effects that generate trophic cascades, so that a direct link can be established between these properties (Figure 1). Dissipative release determines the qualitative nature of the dynamical-structural relationship, while dissipative loss x governs its quantitative expression. When dissipative release is small biomass ratio *B_r_* is proportional to upward response ratio *A*_↑_ (red arrow), with amplification quantified as 1/(1 — *χ*). However, *B_r_* says nothing of the downward response. Conversely, if dissipative release is large, then *B_r_* predicts the downward response, amplified or attenuated by a factor (1 — *χ*)/*χ*. Around the threshold *ξ* ~ 1, downward or upward dynamical effects are entangled in the biomass distribution, so that *B_r_* is partially correlated with both cascade directions (see Figure 3). (b) A within-system relationship between biomass ratio and cascade strength translates into a statistical pattern across ecosystems if they are in similar regimes of *trophic dissipation.* More precisely, any hyperplane of constant *ξ* and *χ* in the larger ecological parameter space leads to a relationship between the shape of the biomass pyramid and trophic cascade strength.

### Box 1. Definition of main concepts.

- **Upward amplification/attenuation:** stronger/weaker response of higher trophic levels relative to lower trophic levels after a perturbation at the bottom of the chain (e.g. nutrient enrichment).
- **Downward amplification/attenuation:** stronger/weaker response of lower trophic levels relative to higher trophic levels after a perturbation at the top of the chain (e.g. fish addition).
- **Self-regulation:** regulatory mechanisms that cause per capita growth rates to depend on the biomass of the species in question (e.g. intraspecific interference, cannibalism or effects from hidden compartments such as pathogens).
- **Trophic dissipation:** general concept representing, at any trophic level, the relative importance of consumer intrinsic loss versus primary productivity (i.e. energy dissipation versus basal influx) in shaping the standing biomass. This concept can be defined formally for a given trophic level, becoming dissipative loss for the consumer, and dissipative release for the resource (Eq. (3)). In every case (including longer and nonlinear food chains), it combines the base magnitude of productivity and consumer losses, and how they are transmitted and modulated by trophic interactions and by self-regulation.

– **Dissipative loss** *χ* in the consumer’s standing biomass, reduction caused by consumer intrinsic loss (from metabolic costs and mortality), divided by gains from primary production.
– **Dissipative release** *ξ* in the ressource’s standing biomass, gains (by prey release) caused by consumer intrinsic loss, divided by gains from primary production.
- **Dynamically consistent chains:** food chains where, in the long-term, the two levels respond in the same direction to nutrient enrichment, and in opposite directions to fish addition.

The key intuition that we explore here is that standing biomasses and trophic cascades both reflect some aspects of the dynamical sensitivity of the food chain. This is obviously true of trophic cascades, but we argue that standing biomasses can also reflect either (i) bottom-up responses of the food chain to primary productivity, (ii) top-down response to consumer losses (e.g. mortality and metabolic costs), or both. Thus, if we can tease apart those contributions we can then infer the chain’s dynamical response to perturbations (Figure 1b). What can and cannot be inferred will clearly be context dependent: for instance, when the rate of consumer intrinsic losses are negligible compared to primary productivity (as in Barbier & Loreau, 2019), the biomass distribution is entirely driven by bottom-up propagation of primary productivity, and cannot tell us whether response to a top-down perturbation (e.g. predator removal) would be weak or strong. Conversely, in some models such as “exploitation ecosystems” (Oksanen *et al*., 1981), standing biomasses are driven by top-down effects, with primary productivity only controlling the number of trophic levels allowed in the food chain.

We reveal a synthetic mechanism that controls the relative contribution of top-down and bottom-up dynamical effects in the biomass pyramid, and thus determines which causal relationships exist, in a given system, between cascades and biomass pyramids. We call this mechanism *trophic dissipation* as it encodes the net effects of consumer intrinsic losses on standing biomasses (Box 1). In particular, this means that primary productivity, trophic interactions or self-regulation (e.g. intra-guild competition) affect the relationship between cascades and pyramids as much as they affect trophic dissipation (Box 1, Figure 1).

Using data from 31 mesocosm experiments of pelagic food webs, which monitored standing biomass and long-term responses to top-down and bottom-up perturbations, we then ask when causal relationships within one ecosystem can translate into statistical patterns across ecosystems (Figure 2b). Combining theory and data, we find that the experimental systems demonstrate two types of relationships between biomass and cascades, i.e. correlations driven by common factors, and direct causation (Figure 1). We conclude that understanding the entangled dependencies between biomass distribution and dynamical responses is essential to properly interpret the patterns observed in natural communities, and enhance our capacity to predict the long-term effects of perturbations.

## 2 Theoretical framework: within-system relationships

We build on the work presented in (Barbier & Loreau, 2019) to develop a theoretical framework spanning the various dynamical regimes previously studied in food chains (e.g. resource and consumer control). Our goal is to determine what dynamical signatures ought to be present, or not, in the standing biomass pyramid of a food chain; and how these signatures can be used to predict the strength of trophic cascades. We complement the framework of (Barbier & Loreau, 2019) by explicitly accounting for consumer intrinsic rate of biomass loss (e.g. due to mortality or metabolic costs). For simplicity, and to match the experimental systems analysed in the next section, we focus on a two-level system, and further assume Lotka-Volterra interactions (i.e. type-I functional responses). However, the theoretical framework presented has no restrictions on the length of the food chain, and can be extended to different functional responses (see Appendix S1 for multi-level and nonlinear chain results). Here, long-term responses to perturbations are understood as shifts in equilibrium biomasses. Although empirical systems are never perfectly stationary, equilibrium analysis of such dynamical models have proven successful in quantifying species interactions and responses even in data with significant temporal variance (*Maynard et al.,* 2019; Barbier *et al*., 2020).

Let *B_i_, i* = 1, 2 be the biomass density of primary producers and consumers, respectively. We model the growth rate of species biomass as

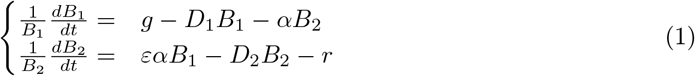

where *g* > 0 denotes the intrinsic biomass production rate of primary producers, and *r* > 0 the intrinsic biomass losses of consumers. The per-capita attack rate of consumers is denoted as *α* > 0 (a rate per unit of biomass density). In the same units we denote by *D_i_ i* = 1, 2 the density-dependent rate of biomass loss at both trophic levels (i.e., self-regulation). Finally, 0 < *ε* < 1 is the efficiency of productivity transfer from producers to consumers. There are two units: time and biomass density. Here we are interested only in non dimensional quantities, such as biomass ratio *B_r_* = *B*_2_/*B*_1_ and response ratios

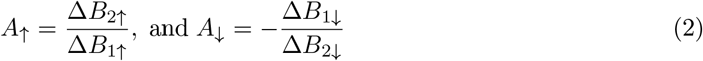

where *A*_↑_ reflects the biomass change of level 2 relative to that of level 1 following a perturbation at the bottom of the chain, while *A*_↓_ represents the biomass change of level 1 relative to that of level 2 following a perturbation at the top. Regardless of the dynamical model considered, if perturbations are not severe enough to elicit significant non linear responses, the use of response ratios as measures of response to perturbations eliminates potential differences in the magnitude of the perturbations performed across systems.

Solving for the equilibrium of a two-species Lotka-Volterra system (see Appendix S2), we find that the biomass ratio *B_r_* can be expressed as

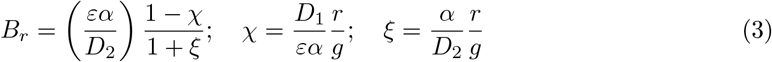

In this expression, we have isolated two terms, *χ* and *ξ*, that express the relative effect of consumer intrinsic losses *r* and primary productivity *g* on the standing biomasses of both levels. Similar terms appear in more complex models (e.g. longer food chains and nonlinear dynamics, Appendix S1).

*χ* is the ratio of predator biomass losses (at equilibrium) due to predators’ intrinsic losses *r*, over gains from consumption of primary productivity *g* (Appendix S1). We call *χ dissipative loss,* as it represents a decrease from the maximum predator biomass allowed by primary productivity. On the other hand, *ξ* is the ratio of biomass gains of the prey due to predators’ intrinsic losses, over the contribution to standing biomass of productivity. We call *ξ dissipative release,* as it reflects the relative importance of prey release caused by consumer losses. Both notions – which are not independent from one another-relate to what call *trophic dissipation*; seen from either the consumer’s (*χ*) or the producer’s (*ξ*) perspective (Box 1).

We now explain how static properties – *B_r_*, the prefactor *εα*/*D*_2_ in Eq. (3), *χ* and *ξ* - can be associated with dynamical responses to bottom-up and top-down perturbations. To see this, consider a perturbation Δ*g* of primary productivity, such as a nutrient enrichment treatment. The changes in biomass of trophic level 1 and trophic level 2 read, respectively

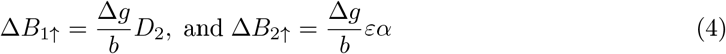

where *b* = *D*_1_*D*_2_ + *εα*^2^ (*b* plays no role in what follows). Similarly, changes in biomass due to a change in consumer mortality Δ*r* (e.g. caused by fish addition), are

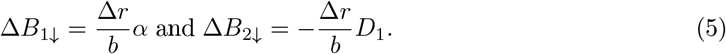

Thus, bottom-up and top-down responses read, respectively

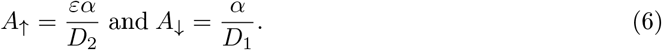

We recognize in *A*_↑_ the prefactor of the r.h.s of Eq. (3). Furthermore, by comparing (6) with the expressions of *ξ* and *χ* in (3) we can deduce that

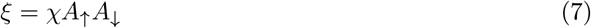

This expression is not just an algebraic manipulation. It reveals the dynamical mechanism that allows us to move from the predator to the prey’s perspective of trophic dissipation. This mechanism is represented by the product *A*_↑_*A*_↓_ of upwards and downwards growth propagation, which measures the strength of the dynamical feedback loop in the food chain (denoted λ in Barbier & Loreau, 2019).

With Eqs. (3, 7) we now have all the ingredients to propose a formal relationship between biomass ratio, response ratio and trophic dissipation (see Appendix S2 for details and generalizations)

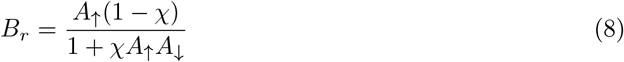

This expression makes it clear that three different factors determine the predator-prey biomass ratio: the two dynamical response ratios *A*_↑_ and *A*_↓_, and the additional effects of growth and losses that enter in the dissipative loss *χ*.

The value of dissipative release *ξ* (7) determines three main qualitative regimes of Eq. (8), as shown in Figure 2 and 3. At the extremes, only upward or downward cascades are related to biomasses:

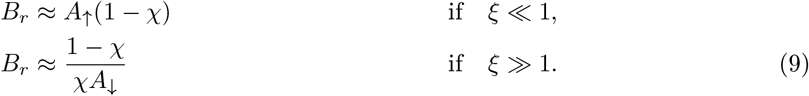

**Figure 3:**
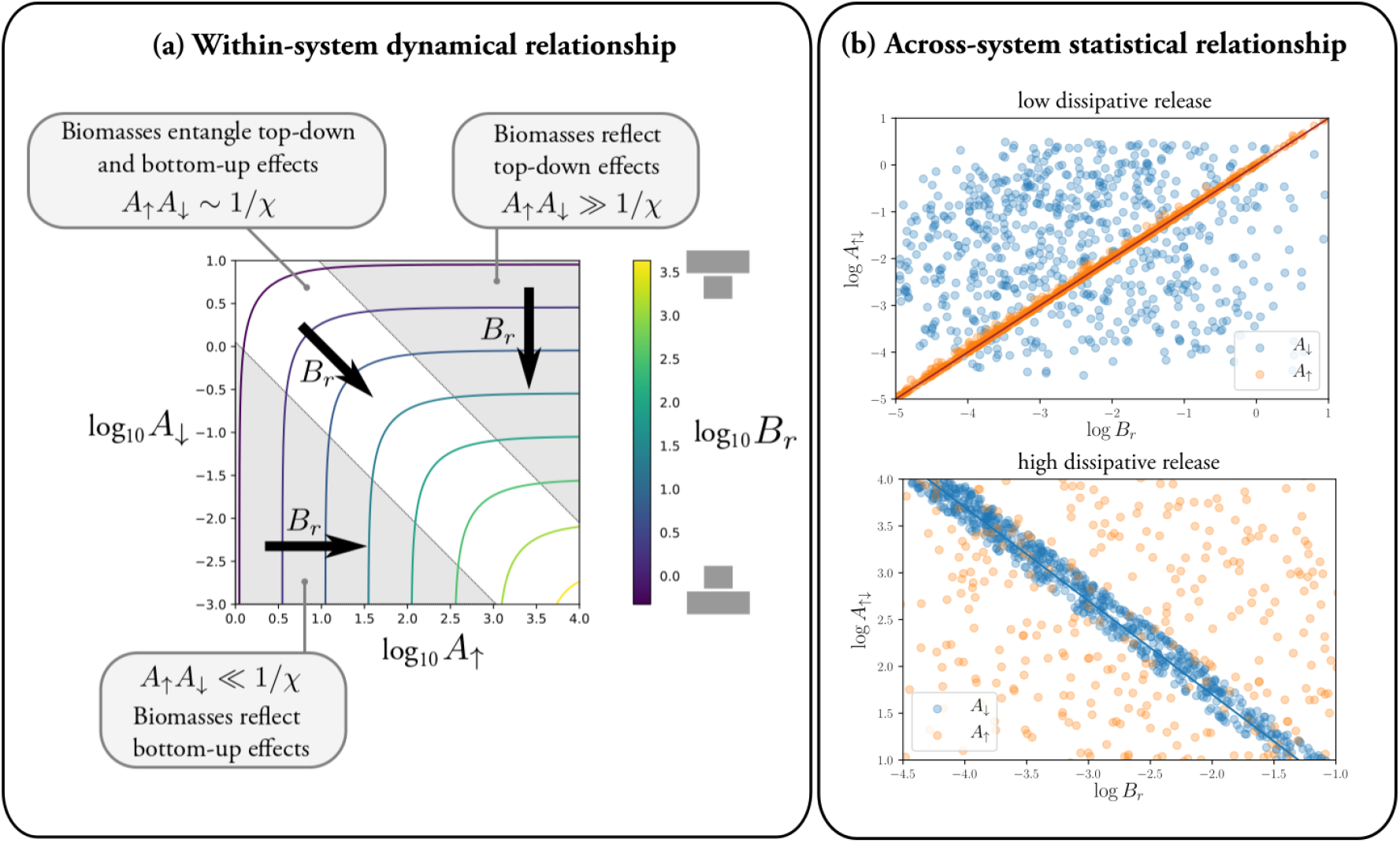
The relationship between biomass distribution across trophic levels and response to perturbations, given dissipative loss *χ* (Box 1). (a) Within a system, if we vary responses *A*_↑_ and *A*_↓_, the threshold 1/*χ* determines the importance of bottom-up and top-down effects. When *A*_↑_*A*_↓_ ≪ 1/*χ* (corresponding to *ξ* ≪ 1 in Figure 2), the biomass distribution only reflects bottom-up effects, as shown by the vertical isolines. Above that threshold, *B_r_* only reflects top-down effects (horizontal isolines), and close to the threshold, it entangles information about both bottom-up and top-down effects. (b) An across-system statistical relationship emerges when there is small variation in *χ*. Simulations of the model (1) (for the range of parameters described in the main text) showing two limit cases of Eq. (8). For low dissipation (here we selected realizations where *ξ* < 10^-1^, and *χ* ∈ (0, 0.4)), upward response ratio *A*_↑_ shows a proportional relationship with biomass ratio (the solid line has slope 1 in log-log scale) while there is no correlation between *B_r_* and downward response ratio *A*_↓_. For large dissipation (here *ξ* > 10 and *χ* ∈ (0.6, 0.8)), we observe the opposite trend: downward response ratio *A*_↓_ is inversely proportional to biomass ratio (solid line has slope −1 on log-log scales) while there is no correlation with *A*_↑_.

For intermediate values *ξ* ~ 1, the contributions of upward and downward responses in the biomass ratio are entangled in Eq. (8). They cannot be separated (and thus up- or downwards cascade strength cannot be predicted from *B_r_*) without knowing the precise value of both *ξ* and *χ*.

We see that *ξ* determines the qualitative regime in which the system is in, while *χ* controls the slope of the relationship within each regime. If *ξ* ≪ 1, then *A*_↑_ ≈ *B_r_*/(1 — *χ*), the biomass distribution is entirely shaped by upward effects. Upward perturbations either follow proportional response if *χ* ≈ 0, as in (Barbier & Loreau, 2019), or are amplified. On the other hand, the biomass ratio contains no a priori information about downward responses *A*_↓_ (Figure 3a). In the opposite limit of large *ξ*, *A*_↓_ will be inversely proportional to B_r_, but with a prefactor (1 — *χ*)/*χ* that can lead to downward cascade amplification if *χ* < 1/2 and attenuation if *χ* > 1/2 (Figure 3b). Proportional response is thus unlikely for top-down effects, as it requires exactly 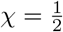. If consumer self-regulation is very weak or zero, we find *A*_↑_, *ξ* → ∞ and only the top-down driven regime is observed, as in classical models (e.g. Oksanen *et al*., 1981; Rossberg *et al*., 2019) that have been used to showcase counter-intuitive dynamical effects in food webs.

## 3 Empirical analyses

### 3.1 Data collection and formatting

To analyse whether causal relationships within one ecosystem translate into statistical patterns across ecosystems, we selected from the literature 21 independent studies reporting on 31 mesocosm experiments of pelagic food webs that performed long-term bottom-up and top-down treatments (see Appendix S2; Hulot *et al*. (2014)). All studies investigated community responses to nutrient enrichment, fish addition, and both perturbations together. They reported species biomass before and after the perturbation. Detailed information about the studies can be found in Appendix S2. These studies allow us to illustrate how our framework can be used to interpret empirical patterns. Even though those patterns might be specific to the pelagic setting of the experiments, we stress that our framework has no restriction on the type of system considered (e.g. marine or terrestrial).

The pelagic communities considered contain different paths of energy transmission across trophic levels (Figure 4a). To properly study biomass pyramids and trophic dynamics, it is fundamental to identify paths of energy that adequately behave as food chains (Barbier & Loreau, 2019). We define as *dynamically consistent chains* those where prey and predators respond in the same direction to bottom-up perturbations, and in opposite directions to top-down perturbations (Box 1). We thus decomposed empirical food webs into all the possible chains and analysed them independently (Figure 4a; Appendix S2). To ensure that the decomposition of the food webs into multiple chains do not bias our analyses we included study system as a random effect in our statistical models (see below).

**Figure 4:**
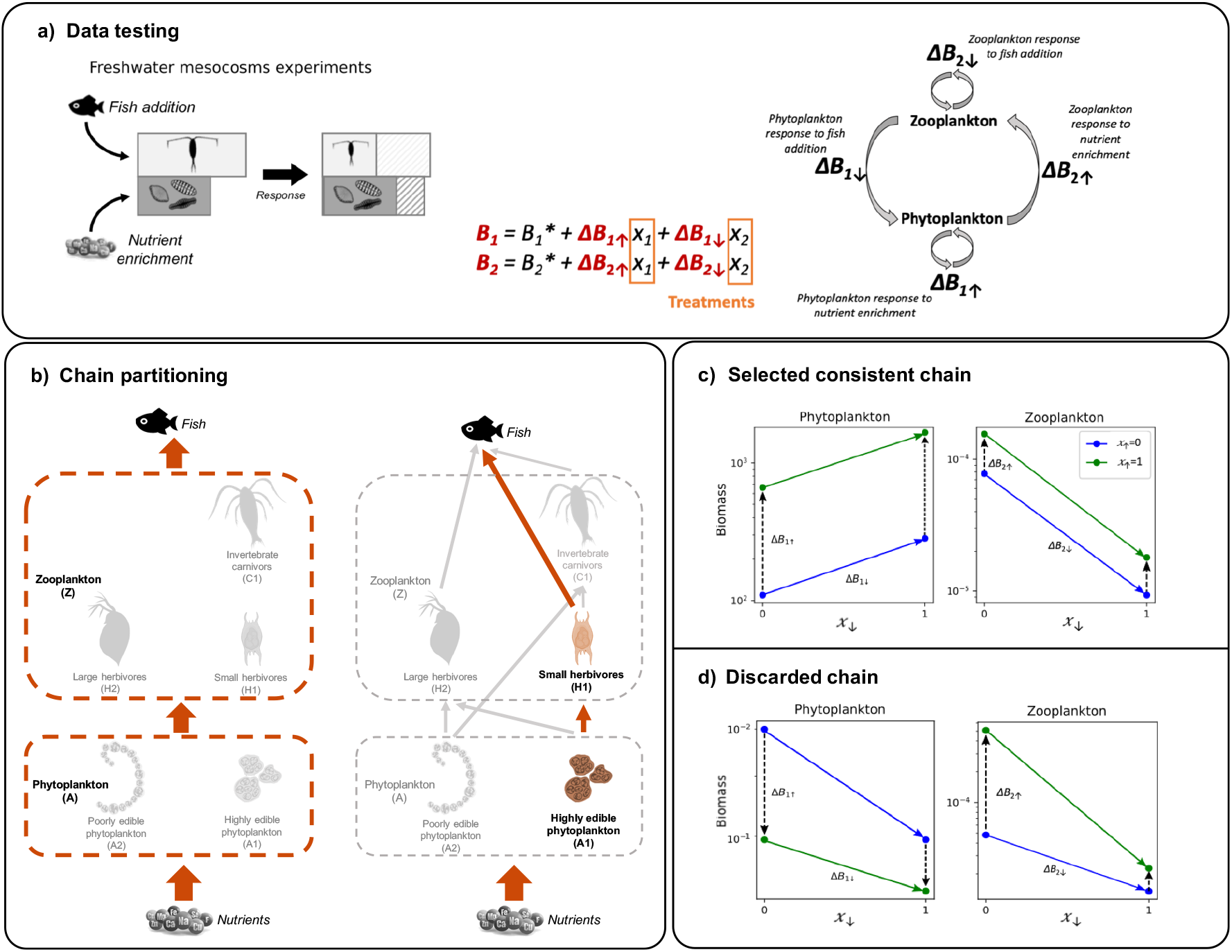
Data analyses illustration. (a) We used data from 31 mesocosm experiments of pelagic food webs that monitored both biomass distributions and long-term responses to nutrient enrichment and fish addition, and estimated the biomass response of two trophic levels to evaluate whether we could detect any of the possible patterns in a natural system. (b) Multiple chains can be identified in these communities. We analysed all possible chains to ensure we capture the dynamics of a *consistent chain* and discarded those chains that did not follow the expected dynamics. Two examples are provided. On the left, all zooplankton species are aggregated into trophic level 2 and all phytoplankton species are aggregated into trophic level 1 (chain: Z-A). On the right, the chain analysed is composed by small herbivores (trophic level 2) consuming highly edible phytoplankton (trophic level 1) (chain: H1-A1). All chains analysed are: Z-A (31), C1-A2 (17), C1-H1 (22), H1-A1 (21), H2-A2 (14), H2-A1 (14), where the numbers in brackets indicate the number of experiments that contained information for each chain type. (c and d) Examples of response evaluation to determine if the chains considered behave as *dynamically consistent chains.* The evaluation was based on the response to both perturbations (i.e., nutrient enrichment and fish addition) of each trophic level. In dynamically consistent chains, we expect the two trophic levels to respond in the same direction to nutrient enrichment, and in opposite directions to fish addition. In (c) we illustrate a case where the system behaved as a dynamically consistent chain. Both trophic levels respond positively to nutrient enrichment (dashed arrows point towards nutrient enrichment), shown by the increase in biomass from blue (no nutrient treatment, *x*_↑_ = 0) to green lines (nutrient enrichment, *x*_↑_ = 1), and they show opposite responses to fish addition (coloured arrows point towards fish addition). Phytoplankton biomass increases from *x*_↓_ = 0 (no fish addition) to *x*_↓_ = 1 (fish addition), while zooplankton biomass decreases with fish addition. In (d) we show a case where the system do not behave as a consistent chain. First, phytoplankton biomass decreases with nutrient enrichment while zooplankton biomass increases. Second, both trophic level respond negatively to fish addition. Additionally, we can also observe the filtering of the responses based on the associated estimated error. While in (c) we observe similar slopes between the green and blue lines (i.e., low error) in (d) the error associated to the estimation of the responses is large, shown by the differences in the slope between green and blue lines.

### 3.2 Data analyses

The third trophic level (fish) was viewed as a perturbation treatment in the experiments considered (only presence/absence reported). Therefore, we characterized the biomass distribution only for the first two trophic levels (phytoplankton and zooplankton), using the ratio

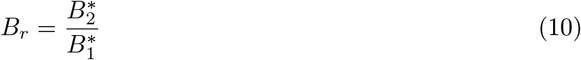

where 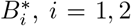 corresponds to the non-perturbed biomass of trophic level 1 (phytoplankton) and level 2 (zooplankton), respectively. *B_r_* quantifies the bottom- or top-heaviness of the biomass pyramid. In particular, if *B_r_* > 1, the pyramid is inverted.

The experiments were designed to observe long-term responses to bottom-up (i.e. nutrient enrichment) and top-down (i.e. fish addition) perturbations (6-12 months). Therefore, we considered the biomasses reported as stationary. To remove the dependency on (unknown) perturbation intensity, we further based our analysis on the expectation of a linear response of biomasses to the two treatments (perturbation intensity is removed by only considering response ratios).

For each level *i* = 1, 2, we estimated the bottom-up change in biomass Δ*B*_*i*↑_ due to nutrient enrichment, and the top-down effect Δ*B*_*i*↓_ due to fish addition. We measured these changes comparing the perturbed biomasses with the unperturbed value 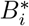. But since there was also a cross-treatment with both enrichment and fish, we fitted a multilinear model

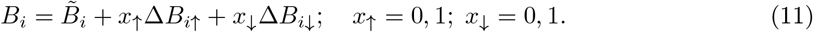

where the binary treatment variable *x*_↑_ = 0 or 1 represents the absence or presence of nutrient enrichment, and *x*_↓_ =0 or 1 represents the absence or presence of fish. We thus identified the coefficients Δ*B*_*i*↑_ and Δ*B*_*i*↓_ as the slopes of the multilinear response, with an intercept 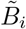.

The goodness-of-fit informed us of whether these estimates were robust: a large fitting error indicates significant interaction between the two treatments. For instance, nutrient enrichment is realized both without and with fish addition. The associated error on response Δ*B*_*i*↑_ is large when the two measurements are incompatible (Figure 4b and c). This can hint either to measurement error, non-stationary behaviour, or strong nonlinearities, which we do not consider here.

We therefore selected the fitted coefficients estimated with low errors (i.e., estimated error < coefficient value, see Figure 4b and c). From selected coefficients, we deduced as in Eq. (2), upward (*A*_↑_) and downward (*A*_↓_) response ratios, where *A*_↑_ measures the biomass change of level 2 relative to that of level 1 following nutrient enrichment, while *A*_↓_ represents the biomass change of level 1 relative to that of level 2 following fish addition.

In addition to the filtering for coefficients estimated with low errors, we only selected the *dynamically consistent chains* (Box 1). Discarded response ratios can indicate other dynamical behaviors (such as competition) that are not consistent with a simple food chain. We verified using Kolmogorov-Smirnov tests that both filters (i.e. low error and expected response direction) did not bias the distribution of the responses considered (Appendix S2).

To analyse the across-system statistical relationship between biomass distribution and trophic cascades, we performed linear mixed effects models including *B_r_* as fixed effects, study system as random effects and *A*_↑_ and *A*_↓_, as response variables. For *A*_↑_ we found that the variance explained by the study system was 0.19, therefore, we included study system as a random effect. However, we found no differences between the model including the study system as random effect (AIC=57.9) and the model including only *B_r_* as fixed effect (AIC=57.0). For *A*_↓_ the variance associated to the study system was 0.002, and the model including only *B_r_* as a fixed effect was better (AIC=58.3; linear mixed model with study system as random effect AIC=60.3). Thus, the results presented below correspond to the analyses were only *B_r_* was included as a fixed effect in the model.

## 4 Across-systems statistical relationship: empirical results

We analyzed 120 potential chains, defined by all the possible paths of energy flux across trophic levels in the experiments considered (i.e. Z-A (31), C1-A2 (17), C1-H1 (22), H1-A1 (21), H2-A2 (14), H2-A1 (14)). We found 68 responses to perturbations that were consistent with that of a dynamical food chain. That is, 33 of the considered paths showed a chain-like behaviour of similar response direction between predator and prey biomass for upward cascades, i.e. *A*_↑_ > 0, while 35 chains showed opposite response direction between predator and prey biomass for downwards cascades, i.e. *A*_↓_ > 0 (Table S1.2 in Appendix S2).

Paths where all zooplankton species and all phytoplankton species were grouped together to form two trophic levels (chain Z-A in Figure 4a) showed the highest percentage of chain-like responses across all experiments, together with the chain composed by small herbivores (H1) consuming highly edible phytoplankton (A1). The estimated responses to perturbations had low fit significance or wrong signs in approximately 60% of the instances (61% for Δ*B*_1↑_, 68% for Δ*B*_2↑_, 58% for Δ*B*_1↓_ and 60% for Δ*B*_2↓_, cf. Figure S2.1 and S2.2). For instance, we observed zooplankton biomass increasing after fish addition, or phytoplankton biomass decreasing due to nutrient enrichment.

The biomass distribution was bottom-heavy (i.e., *B_r_* < 1) in most selected paths (Figure 5). In the selected experimental systems, *B_r_* was positively correlated with *A*_↑_. In log scale, the slope of this relationship was close to one with an intercept close to zero, indicating that the responses to nutrient addition of each trophic level were proportional to their biomass, showing no further effect of trophic interactions in amplifying the upwards response.

**Figure 5:**
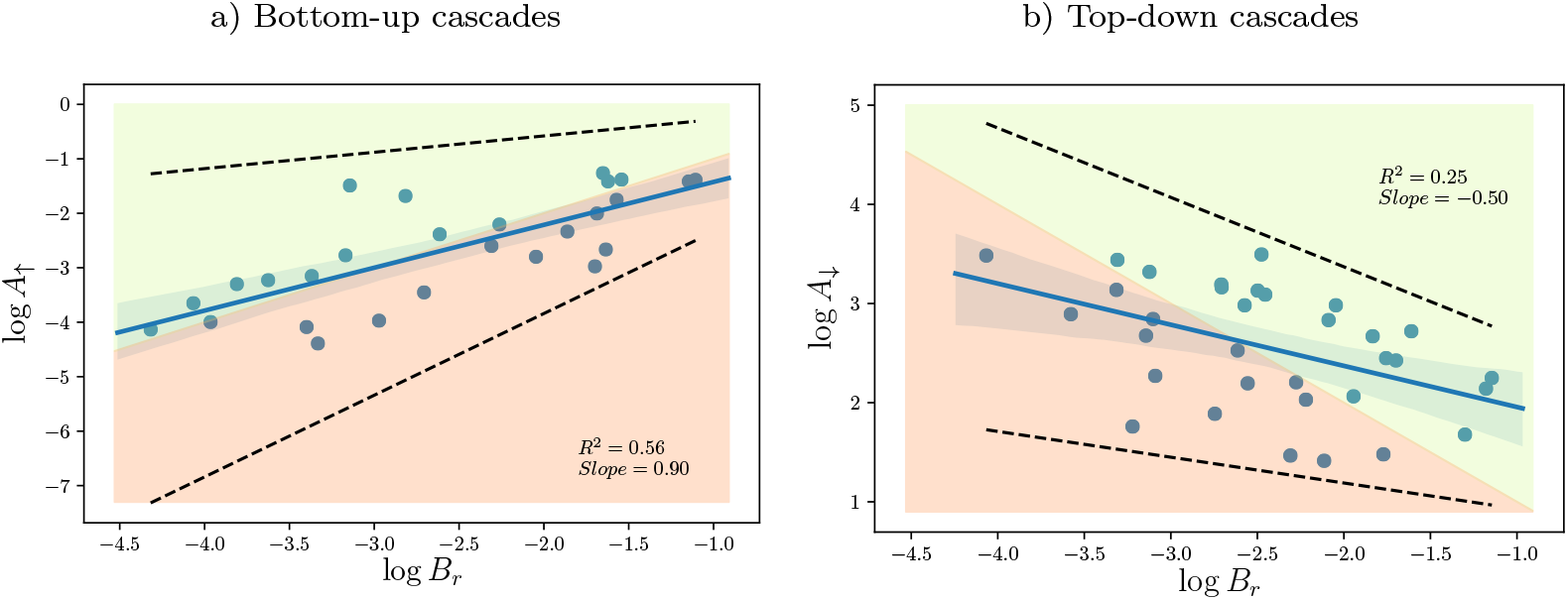
Empirical relationship between biomass ratio and response ratios. (a) Relationship between biomass ratio and bottom-up response ratio 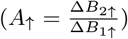 due to nutrient enrichment. In (b) relationship between biomass ratio and top-down response ratio 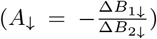 due to fish addition. Each point represents a food chain in our analyses. Blue lines represent the linear regression and shaded areas represent 95% confidence intervals. Dashed lines represent the limits of the empirical pattern used in the simulation analyses to filter the prior distribution of parameters. Coloured regions represent the simple expectation scenario where a trophic level’s response to a perturbation is proportional to its biomass. In the green region perturbations are amplified through the chain while in the orange perturbations are attenuated. Thus, the interface between the two regions represents the 1:1 line where the response of each trophic level is proportional to their biomass. Notice that patterns shown in (a) and (b) are excluding the outliers, which results into an exclusion of the extremes of the biomass ratio distribution (See Figure S2.3 in Appendix S3 for results including the outliers).

In contrast, *B_r_* was negatively correlated with *A*_↓_ with a slope of −0.5 in log scale (Figure 5b). Yet, as we discuss below, this trend may be spurious, if top-down effects are not causally involved in shaping the biomass ratios.

## 5 Comparing theory and data: Approximate Bayesian Computation (ABC)

We can use our theoretical results to understand the empirical patterns and underlying mechanisms. We used ABC to identify parameter ranges consistent with the data (Beaumont, 2010). The idea of ABC is to leverage empirical patterns to constrain the distribution of simulation parameters. In this way, we can use across-system patterns to infer within-system dynamics. We generated predator-prey systems with different parameter values drawn from broad prior distributions, and computed their equilibrium biomass ratios *B_r_* and top-down (*A*_↓_) and bottom-up response ratios (*A*_↑_). We then retained only model realizations for which (i) the two species coexisted and (ii) the relationship between *B_r_* and *A*_↑_ and between *B_r_* and *A*_↓_ were consistent with those observed in our data analyses (Figure S3.1). We retained 1500 model realizations, and used the corresponding parameter values to compute a joint posterior distribution for all model parameters. We thus simulated systems in which the theoretical within-system relationship holds exactly, and infer how the variation of key ecological parameters can explain the empirical relationship across systems. All parameters were drawn uniformly on a log scale, over a span of several orders of magnitude: biomass conversion efficiency *ε* ∈ [10^-2^,10^-1^]; relative attack rate 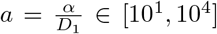; relative self-regulation 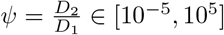; and relative intrinsic rate 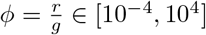.

We note that, although we did not constrain the model parameters to match the empirical biomass ratios (but only to follow the relationship of biomass and response ratios), the selected simulated data points displayed a remarkably similar distribution of biomass ratios (Figure S3.1). Once filtered to match empirical patterns, the distribution of dissipative release *ξ* (i.e. prey release due to dissipation) shifted towards small values (c. 67% had *ξ* < 0.1 cf. Figure 6a). This suggests that for two thirds of systems (c. 67%), biomass ratios were shaped by bottom-up effects. The empirical trend between biomass ratio and top-down effects was allowed in our model and simulations, yet mainly driven by a weak anticorrelation between upward and downward response (Figure S3.1). Grouping simulations by dissipative release (low, intermediate and high), we found that the three classes presented qualitatively similar patterns with top-down responses (Figure S3.2), with a slight steepening of the slope for systems with strongest dissipative release.

**Figure 6:**
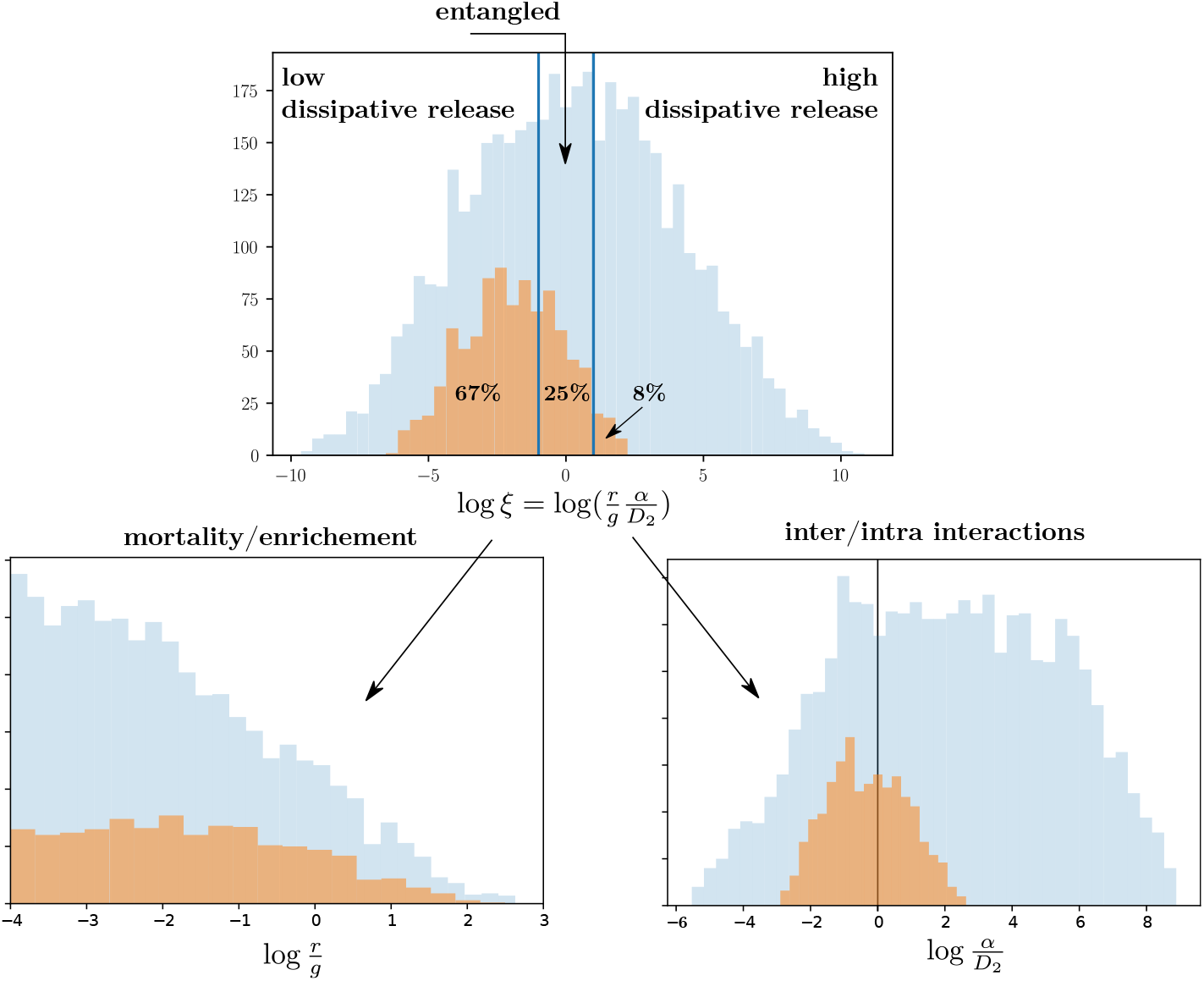
Prior and posterior distributions of model parameters. Blue histograms represent the prior distribution of the parameters introduced into the model (conditioned on coexistence). Orange histograms represent the distribution of the parameters after filtering them to match empirical patterns. Top: distribution of *ξ* = (*r/g*)(*α/D*_2_) (dissipative release) which determines the nature of within-system relationship between biomass ratio and upwards and downwards response ratios. Dissipative release is the product of two non dimensional ratios. Lower left: distribution of the ratio of predator intrinsic loss to primary productivity *ϕ* = *r/g,* Lower right: distribution of the ratio *α/D*_2_, which represents the relative strength of trophic interactions and predator self-regulation. We see that the posterior distribution is centered close to 0 in log scale, suggesting that self-regulation rates are comparable to predation rates.

In terms of model parameters, dissipative release reads *r/g* × *α/D*_2_. The distribution of *ϕ* = *r/g* remained broad after filtering by the empirical patterns (Figure 6b). On the other hand, the ratio of attack rate and consumer self-regulation *α/D*_2_ was more constrained by the empirical patterns: its posterior distribution shows a unimodal distribution centered around 1 (Figure 6c). Thus, the ratio *r/g* may be driving the across-system variation in dissipative release.

## 6 Discussion

Trophic cascades are widespread in nature (Pace *et al*., 1999; Shurin *et al*., 2002; Estes *et al.,* 2011), where changes in the biomass of a given trophic level ripple along the food chain through both direct and indirect biotic interactions, ultimately altering ecosystem structure and functioning (Paine, 1969; Carpenter *et al*., 1985; Polis *et al*., 2000; Duffy, 2002; Estes *et al*., 2011).

Understanding the prevalence and strength of trophic cascades has occupied a large part of the food chain literature (Shurin & Seabloom, 2005; Borer *et al*., 2005; Heath *et al*., 2014; Barbier & Loreau, 2019; Sentis *et al*., 2020). Here, rather than investigating the drivers of trophic cascades, we explored the nature and drivers of *the relationship* between cascade strength and the more empirically accessible notion of biomass pyramids (Elton, 1927)(Figure 1). We do not ask why cascades are weak or strong, but rather why their strength can or cannot be predicted from the biomass pyramid.

Although previous studies have discussed whether the static properties of the food chain, such as its biomass pyramid or its size spectra (the distribution of organism body size) can hint at the underlying dynamics (McCauley *et al*., 2018; Barbier & Loreau, 2019; Rossberg *et al*., 2019), a systematic understanding of the link between static and dynamical properties of food chains is still lacking. Our contribution to bridge this gap is to provide a general and synthetic description of the relationship between the strength of trophic cascades and the shape of biomass pyramids.

### 6.1 Summary of results

Using both theory and data from pelagic experiments, we asked under which conditions the strength of trophic cascades can have a signature in the biomass pyramid.

Our results are based on the realisation that cascading responses to perturbations *and* the biomass pyramid, both reflect some aspect of the dynamical sensitivity of a food chain (Figure 1b). Indeed, the standing biomass distribution is the product of a bottom-up dynamical response to primary productivity, as well as a top-down response to consumer intrinsic losses. Our approach was to determine the relative contributions of those bottom-up and top-down dynamical processes, to then predict the relationship (or lack thereof) between the shape of the biomass pyramid and either bottom-up, top-down cascades or a combination of both. We stress that this question is not equivalent to asking whether the system is bottom-up or top-down controlled (as in e.g. Kokkonen *et al*., 2019), but whether bottom-up or top-down dynamical effects, relevant to predict trophic cascades, can be seen in the biomass distribution.

We showed that the contribution of bottom-up and top-down dynamical effects in the biomass pyramid is determined by *trophic dissipation*, which can be seen from either the consumer’s or the resource’s perspective (dissipative loss and release, respectively -see Box 1), and gives us the key combinations of primary productivity, predator losses, self-regulation and attack rates that determine the relationship between cascades and biomass pyramids. These perspectives are complementary: *ξ* determines three qualitative scenarios, while *χ* controls the slope of the relationship. Low dissipative release, *ξ* ≪ 1 implies that standing biomasses only reflect bottom-up dynamical effects, which are the ones that determine the strength of bottom-up cascades. Conversely, *ξ* ≫ 1 indicates that only top-down dynamical effects (which determine top-down cascades) shape the equilibrium state. For intermediate values, *ξ* ~ 1, bottom-up and top-down effects become entangled, so that the biomass pyramid does not provide precise information about the strength of either bottom-up or top-down cascades, but rather a combination of the two. We noted that, by construction, many classic food chain models such as exploitation ecosystems (Oksanen *et al*., 1981) are in the singular top-down limit *ξ* → ∞.

The crucial distinction between asking whether cascades are strong and whether they are causally reflected in the biomass distribution, is clearly demonstrated here: top-down cascades can be predicted from the shape of the biomass pyramid if natural predator losses are sufficiently high (*ξ* > 1), but the larger these losses, the weaker the cascades, since the predator population is more depleted, with less potential for further prey release from perturbations.

The existence of a pattern across multiple systems is contingent on low variation of dissipative losses *χ*, while the pattern’s shape depends on the magnitude of dissipative release *ξ* (Figure 3). Using Approximate Bayesian Computation (ABC), we estimated the distribution of dissipative release compatible with the observed empirical pattern. We found that experimental data was largely compatible with the bottom-up (low dissipative release, 67%) or entangled (25%) scenarios. Only 8% of systems had high enough dissipative release for top-down effects to determine the biomass ratio (Figure 6). Nonetheless, we did observe a trend between biomass ratios and topdown effects (Figure 5b). This relationship mainly arose from latent parameters that create an anticorrelation between *A*_↓_ and *A*_↑_, and subsequently between *A*_↓_ and *B_r_* (Appendix S3). Only a small contribution to the trend came from the fraction (25% + 8%) of systems where topdown effects play direct, causal role. Even if we observed a similar relationship between downward responses and biomass ratios in all three groups of systems, our theory proposes that those patterns should be interpreted differently; from a spurious correlation for low dissipation release to an actual causation for high dissipation release.

Beyond hinting at which empirical patterns are causal, our framework may be used to shed light on specific ecological parameters. The ABC analysis suggested that intra- and inter-level interactions are of roughly comparable strength in these pelagic experiments, while the balance of primary productivity and consumer intrinsic losses may be the most important axis of variation in dissipative release between experiments.

### 6.2 Limitations

Our theoretical framework was used to analyse data pooled from multiple experimental pelagic communities. The empirical patterns reported and their interpretation might therefore be specific to pelagic systems. However, our theoretical framework is as general as the notion of food-chain itself (and linear response to perturbations). It has no restriction on the type of system considered, and could thus be used to analyse the drivers of the relationship between trophic cascades and biomass pyramids across ecosystem types.

Only 50% of the empirical systems displayed a consistent chain-like response to either bottom-up and top-down perturbations, and very few had both. This is unfortunate, as using information from both types of perturbations would have allowed us to infer relevant ecological parameters (e.g. self-regulation) in specific communities, rather than only trends across systems.

Different mechanisms could explain this high rate of discrepancies, among which omnivory is a prominent one. The presence of omnivorous links in food webs has been well documented (Polis *et al*., 1989; Polis, 1994; Emmerson & Yearsley, 2004; Neutel *et al*., 2007), and is known to obviate cascading effects (Diehl, 1993; Pringle & Hamazaki, 1998; Polis, 1999; McCauley *et al*., 2018). Indeed, many ‘herbivorous’ zooplankton species consume phytoplankton but also a substantial amount of microzooplankton, such as rotifers and ciliates (Sprules & Bowerman, 1988; Gilbert, 1988; Brett *et al*., 1994; Hansson *et al*., 2004), which can explain some of the non-chain-like responses observed. Similarly, compensation between species or functional groups within one trophic level (Gonzalez & Loreau, 2009) could also attenuate or reverse the expected dynamical response (Leibold, 1989; Hunter & Price, 1992; Strong, 1992; McCann *et al*., 1998).

Additionally, we may be unable to correctly identify chain-like dynamics when the response of either level to a perturbation is very small, and indistinguishable from measurement noise. Taking a ratio of responses is necessary to eliminate the confounding role of perturbation intensity, but this ratio becomes poorly defined when either response is close to zero.

We obtained the largest fraction of chain-like responses when considering all zooplankton species and all phytoplankton species as two trophic levels, suggesting that consistent chain dynamics may emerge as a complexity reduction of communities into few interacting levels composed by multiple species or groups of species (Ulanowicz, 1995). This suggests a complementary line of research: identify the conditions under which a food web behaves as a chain, thus allowing our theory to apply.

### 6.3 Influence of primary productivity

A long-standing empirical and theoretical question, is how primary productivity can affect either biomass distribution (McNaughton *et al*., 1989; Del Giorgio *et al*., 1999) or trophic cascade strength (Carpenter *et al*., 1985). Here, we showed how primary productivity can change the *relationship* between biomasses and cascades. Increasing primary productivity lowers the strength of trophic dissipation, leading to biomass pyramids shaped by bottom-up propagation of primary productivity that convey, a priori, no information about top-down cascades. Our ABC approach suggests that the variation in dissipative release across systems was mainly driven by a variation in the ratio of intrinsic rates (consumer mortality over prey productivity) among systems. Thus, variations in primary productivity could be an essential factor explaining the diversity of observed pyramids and dynamical responses in the pelagic systems analysed.

### 6.4 On the importance of self-regulation

Previous research has already pointed at the importance of self-regulatory processes to modulate the strength of bottom-up and top-down effects (McCann *et al*., 1998; Herendeen, 2004; Heath *et al*., 2014). Our theoretical framework elucidates how both resource and consumer self-regulation shape the link between static and dynamic properties in a food chain, as they determine the importance of trophic dissipation in the observed biomasses of consumers and resources.

Very few empirical studies, however, provide reliable direct estimates of consumer self-regulation (Skalski & Gilliam, 2001). Furthermore, effects from species or compartments outside of the considered chain could participate in the observed self-regulation (Loreau, 2010). This would make it an emergent property that is not directly accessible from individual behaviour such as predator interference, but could still be estimated indirectly through our framework.

## Conclusion

Cascading responses to perturbations and the biomass pyramid both reflect some aspects of the dynamical sensitivity of a food chain. Based on this observation, we asked whether the shape of the latter could be used to predict the strength of the former. Our approach was to determine the relative contributions of bottom-up and top-down dynamical effects to the biomass distribution, to then predict the relationship (or lack thereof) between the shape of the biomass pyramid, and the strength of either bottom-up or top-down cascades. We identified trophic dissipation, as the driver of this relationship. Trophic dissipation is the relative contribution of consumer intrinsic losses in the standing biomass distribution. It depends on well-studied ecological parameters, such as primary productivity and attack rates, but also on the less empirically accessible notion of prey and predator self-regulation. We noted that an observed relationship between biomass distribution and upward or downward cascades can reflect a common dynamical cause, but can also arise from correlations driven by a common factor. In the case of a common dynamical cause, we can extrapolate this relationship to make new predictions, within or across ecosystems, without additional knowledge of confounding factors. Our results thus provide criteria for when measurements of static properties can be used as reliable indicators to predict food chains’ dynamical response to perturbations.

## Acknowledgments

We thank Bart Haegeman for helpful discussions and review of previous versions of the manuscript. We also thank Florence Hulot for sharing experimental data. This work was supported by the TULIP Laboratory of Excellence (ANR-10-LABX-41) and by the BIOSTASES Advanced Grant, funded by the European Research Council under the European Union’s Horizon 2020 research and innovation program (666971). JFA was supported by an Irish Research Council Laureate Awards 2017/2018.

## Author contributions

All authors contributed to the design of the study. NG and AA assembled the empirical data and, jointly with MB, performed the empirical analyses. JFA and MB developed the theoretical framework. NG and JFA performed the numerical simulations. NG, MB and JFA wrote the first draft of the manuscript, and all the authors contributed substantially to revisions.

## Supporting Information

### Appensix S1. Theoretical framework

#### Two-level chain

Let *B_i_, i* = 1, 2 be the biomass density of primary producers and consumers, respectively (e.g grams per unit of volume). We denote by *g* > 0 the intrinsic growth rate of primary producers, and *r* > 0 the intrinsic death rate of consumers. The per-capita attack rate of consumers is denoted as *α* > 0 (a rate per unit of biomass density). In the same units we denote by *D_i_ i* = 1,2 the density-dependent rate of biomass loss (or reduction of productivity) at both trophic levels (i.e., self-regulation). Finally, we write 0 < *ε* < 1 the (non dimensional) efficiency of productivity transfer from primary producers to consumers. This leads us to the following expression for the growth rates 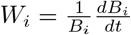 of both populations

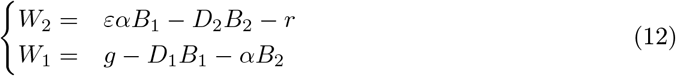

which define a Lotka-Volterra dynamical system (See Appendix S1 for the nonlinear chain approximation). In what follows we are interested only in non dimensional ratios, such as the biomass or response ratio of the two levels. We therefore begin by adimensionalizing the problem. There are two units: time and biomass density. To remove the former we can express time relatively to the rate of growth g of primary producers. We write *τ = gt* (non dimensional time) and, since 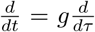, expressing the dynamics in those units amounts to dividing by g all terms of the right hand side of (12). To remove the units of biomass, we can express all densities with respect to the carrying capacity *K*_1_ = *g/D*_1_ of primary producers. We thus define *b_i_* = *B_i_*/*K*_1_ (non dimensional biomass). This lead us to the non dimensional system, denoting 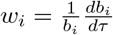 non dimensional rates, we get that

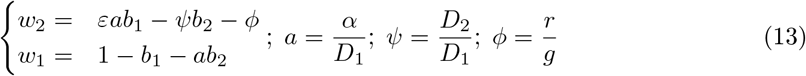

At equilibrium 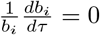 so that (if both species coexist), their respective densities satisfy

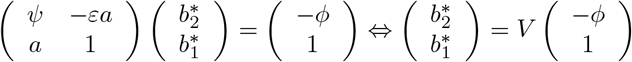

where the matrix *V* is

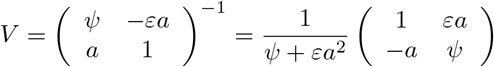

We deduce, in particular, the biomass ratio 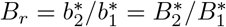 equal to

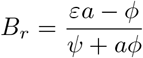

where we can observe that the coexistence condition is that *εa* (consumption) must be larger than *ϕ*. We can also recognize in this expression of *B_r_* the amplification of perturbations. This is because the matrix *V* that gave us the equilibrium densities also gives us their sensitivity to changes in growth rates. Indeed, consider a perturbation of the primary producer in the form of a persistent change *δg* of growth rate. In the non dimensional formulation, this is a perturbation *δw*_1_ = *δg/g* and it leads to the shift of biomass:

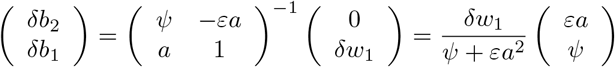

where we recognize the second column of *V*, times *δw*_1_. We can do the same for a perturbation *δw*_2_ of the consumer’s growth rate, which would give us the first column of *V*, times *δw*_2_. Thus

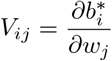

so that bottom-up response ratio (i.e., perturbation applied on resources), defined as 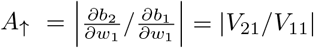 reads

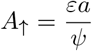

Similarly, if the perturbation is applied to the consumer we get that top-down response ratio 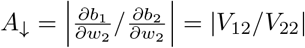 is in fact

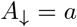

From these notations yields the following expression for the relationship between biomass ratio and trophic cascades:

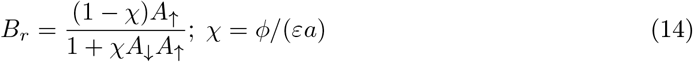

Where *χ* can be interpreted as a measure of dissipative loss, the ratio of biomass losses of the consumer, over gains from primary productivity. If instead we were to define *ρ* = *χA*_↑_ we would have had

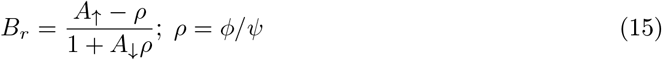

Where 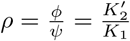 can be interpreted as the ratio of intraspecific characteristic densities: carrying capacity *K*_1_ for the producer, and the characteristic density 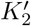 of consumer population where intraspecific competition affects the growth as much as intrinsic loss. In particular, constant *ρ* would relate the biomass ratio to the amplification factors, which signal whether cascades are weak or strong relative to the proportional baseline. We get that:

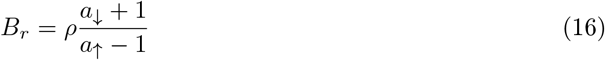

This expression can be misleading, because the amplifications *a*_↑_ = *A*_↑_/*B_r_* and *a*_↓_ = *A*_↓_ × *B_r_*, are not independent of the biomass ratio, which is in turn shaped by *ρ*: in particular, we know from (8) that, if *ρ* = 0, then *a*_↑_ = 1. What this expression shows, however, is the confounding role of *ρ*. If this parameter varies across systems more than either amplification factor, no relationship between *B_r_* and cascade strength will be found. If, however, *ρ* has low variance across ecosystems while *a*_↓_ varies significantly, then we can expect a relationship such as the one observed in Figure 5e.

#### Multi-level chain

The above reasoning lends itself to a generalization to arbitrary number levels. If *α*_*i,i*–1_ is the attack rate of level *i* on the level below, *D_i_* is the strength of intraspecific competition and *r_i_* is consumer intrinsic rate of biomass losses, due to metabolic costs and mortality. As for the the two level system, removing dimensions leads to the system

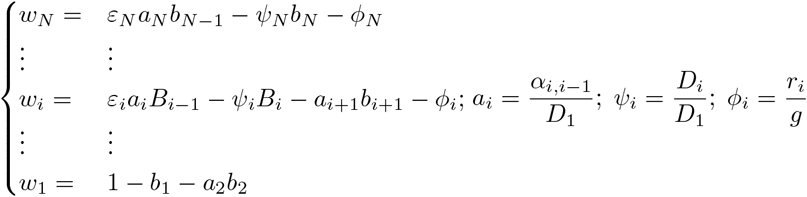

From these we could compute the sensitivity matrix *V* = (*V_ij_*) as previously defined. It is the inverse of the interaction matrix *α* = (*α_i,j_*)

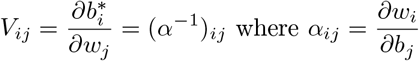

which takes the more explicit form here

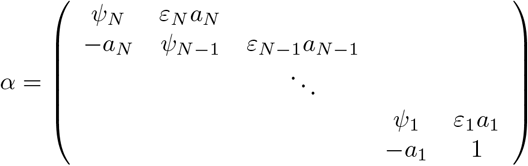

The equilibrium densities (assuming coexistence) are then given by

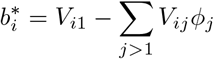

Response ratios now read, if *j > i* (upward response ratio)

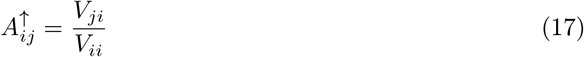

and the same holds if *j < i* (downward response ratio 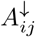). The biomass ratio between top and lowest level reads

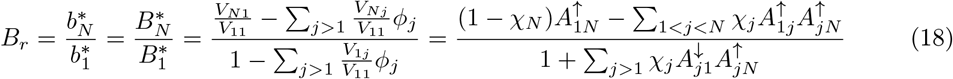

where we defined dissipative loss at level *j*

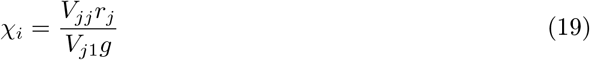

thus showcasing the generality of our approach: top down responses are causally related to top-to-bottom biomass ratio if total dissipative release 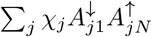 is comparable (or larger) than 1. Note that having many levels is likely to extend the entangled domain, where both upwards and downwards transmission of perturbations shape the biomass distribution.

#### Nonlinear chain

In fact, the above results are not specific to linear growth rates (i.e Lotka-Volterra). For any form of growth rate functions *w_i_* we can still define the interaction matrix 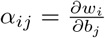 and the write the density independent part of those functions —*ϕ_i_* for consumers 1 for the primary producer (intrinsic growth rate, used to set a time scale). As before 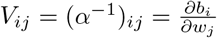 and we can therefore write

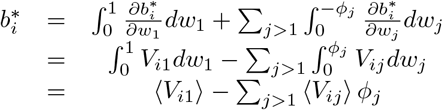

where 〈*V_ij_*〉 is the average sensitivity of species *i* to a change in growth rate of species *j*, as relative loss is varied from 0 to its actual value, *ϕ_j_*. We now have

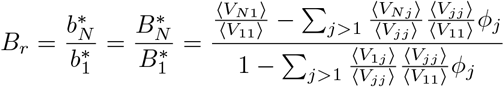

The difference with the linear chain is that now the response ratio of a weak perturbation will be (for instance)

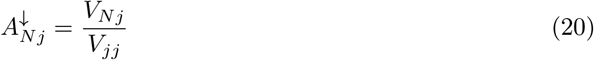

evaluated at the actual conditions defined by the relative intrinsic rates *ϕ_i_*. The response ratio is not exactly equal to the term

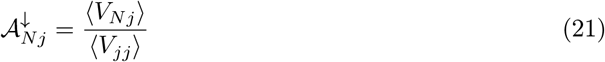

that appears in the above expression of biomass ratio. Nonetheless, unless the growth rates *w_i_* are strongly nonlinear functions of 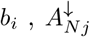 and 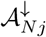 (in particular) can be expected to be strongly correlated. If

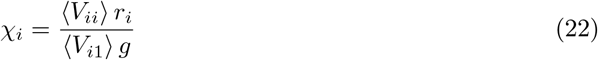

we have that

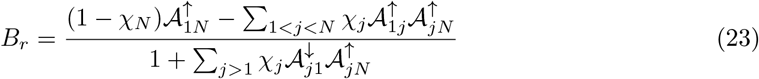

with 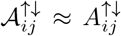. This extension is useful to understand why nonlinearities can allow for some patterns in the data that would be forbidden by a the linear chain model. Indeed, if we go back to the expression of *B_r_* in two level chain (14) we see that *A*_↑_ must be strictly larger than *B_r_*. In the data this is not always the case. Slight non-linearity will remove this strong constraint since it is now 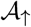 that must be larger than *B_r_* and not *A*_↑_. As we show numerically, only slight non-linearity (in the form of handling time by the consumers) is enough to produce theoretical pattern consistent with the data, as shown in Figure (S1.1).

**Figure S1.1:**
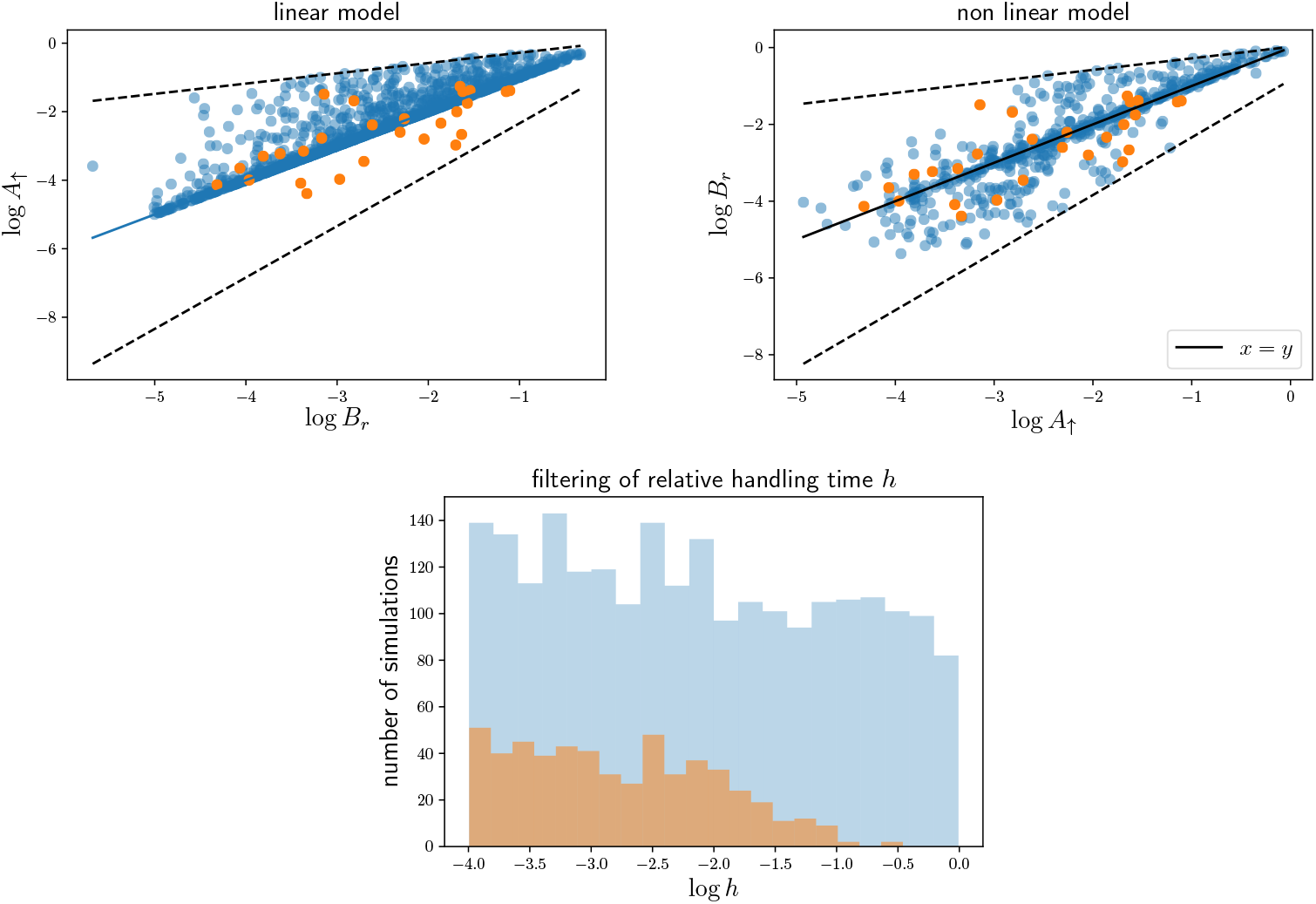
Comparison of simulations and data for linear and non-linear resource-consumer models. In the non-linear model, we use a Holling type II functional response 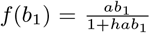 for consumption, with relative handling time *h*. All other parameters are the same, and drawn from the same distributions, as in the linear model described in the main text, and *h* is drawn uniformly on log scales in [10^-4^,10^0^]. Upper row: *A*_↑_ vs *B_r_* for empirical data points (orange) and simulated data points (blue). In the linear model (left), *A*_↑_ cannot be smaller than *B_r_*, yet the data shows points that violate this condition. (right) slight non linearity can relax this constraint. Bottom row: distribution of handling time after filtering by the empirical patterns. We observe that only small values are selected (weak non linearity).

### Appendix S2. Empirical data

**Table S2.1:** Experimental studies used in the analyses. We extracted 31 experimental studies analysing bottom-up and top-down effects in pelagic food chains after press perturbations. Detailed information about the studies used can be found in (Hulot *et al*., 2014).

| Article | N. of experiments |
| --- | --- |
| (Vanni, 1987a,b) | 2 |
| (Van De Bund & Van Donk, 2004) | 2 |
| (Lacroix & Lescher-Moutoue, 1991) | 1 |
| (Fernández-Aláez *et al.*, 2004) | 2 |
| (Stephen *et al.*, 2004) | 2 |
| (Lynch & Shapiro, 1981) | 1 |
| (Menezes *et al.*, 2010) | 1 |
| (Qin & Culver, 1995) | 1 |
| (Hansson *et al.*, 2004) | 1 |
| (Romo *et al.*, 2004) | 2 |
| (Dickman *et al.*, 2008) | 1 |
| (Acuña *et al.*, 2008) | 1 |
| (Beresford & Jones, 2010) | 1 |
| (Drenner *et al.*, 1990, 1989) | 2 |
| (Faafeng *et al.*, 1990) | 2 |
| (Hietala *et al.*, 2004) | 2 |
| (McQueen *et al.*, 1992) | 1 |
| (Proulx *et al.*, 1996) | 2 |
| (Sinistro, 2010) | 1 |
| (Riemann, 1985) | 1 |
| (Mazumder *et al.*, 1990) | 1 |

**Table S2.2:** Total number of experiments analysed per chain type and number of chain-like responses for upwards amplification and downwards amplification for each chain type.

| Chain type | Total number of experiments | Upwards amplification | Downwards amplification |
| --- | --- | --- | --- |
| Z-A | 31 | 10 | 12 |
| C1-A2 | 17 | 4 | 2 |
| C1-H1 | 22 | 5 | 6 |
| H1-A1 | 21 | 7 | 7 |
| H2-A1 | 14 | 3 | 6 |
| H2-A2 | 14 | 4 | 2 |

**Figure S2.1:**
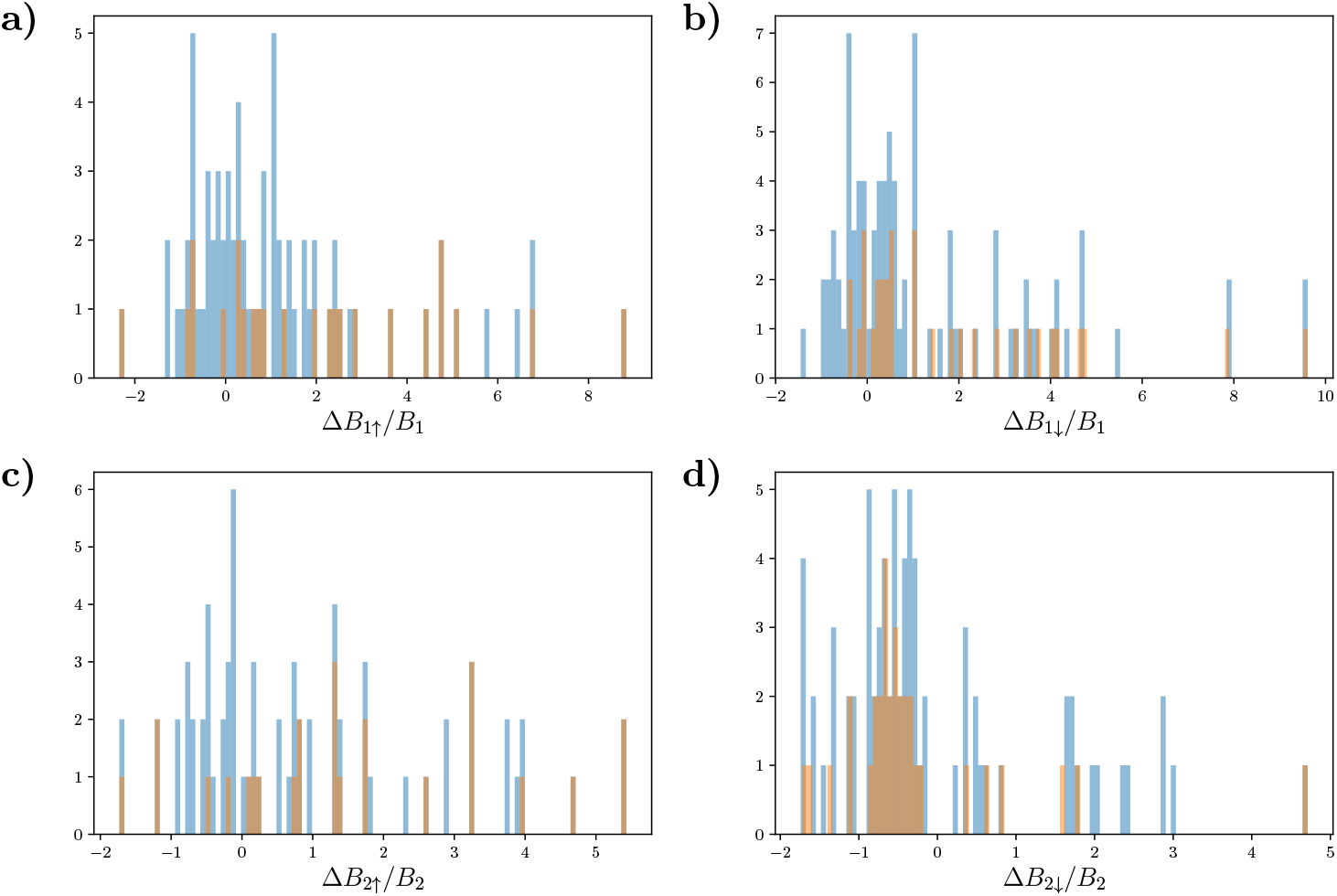
Distributions of the estimated responses to nutrient enrichment (↑) and fish addition (↓) treatments of each trophic level biomass (*B*_1_ and *B*_2_, for trophic level 1 and 2, respectively) relative to their original biomass in experimental mesocosms. Blue distributions represent the estimated responses of all possible chains analysed before filtering them to ensure they behave like dynamically consistent chain and before selecting the ones that presented estimated responses with low error. Orange distributions show the estimated responses of the chains that were identified as dynamically consistent chains. In dynamically consistent chains, we expect the two levels to respond in the same direction to nutrient enrichment, and in opposite directions to fish addition. More concretely, we expect responses of trophic level 1 to nutrient enrichment (a) and to fish addition (b) to be generally positive, together with the biomass responses of trophic level 2 to nutrient enrichment (c). Therefore, most of the chains selected present a positive biomass response to the perturbation. In contrast, biomass responses of trophic level 2 to fish addition (d) are expected to be generally negative. Thus, we observe that very few chains are selected when trophic level 2 presents a positive response to fish addition. In the following lines we present the statistical results from the comparison between the filtered and nonfiltered distributions using Kolmogorov-smirnov tests for each response. If the K-S statistic is small or the p-value is high, then we cannot reject the hypothesis that the two distributions are the same. a) statistic = 0.253, p-value = 0.167; b) statistic = 0.197, p-value = 0.271; c) statistic = 0.251, p-value = 0.165; d) statistic = 0.17, p-value = 0.476

**Figure S2.2:**
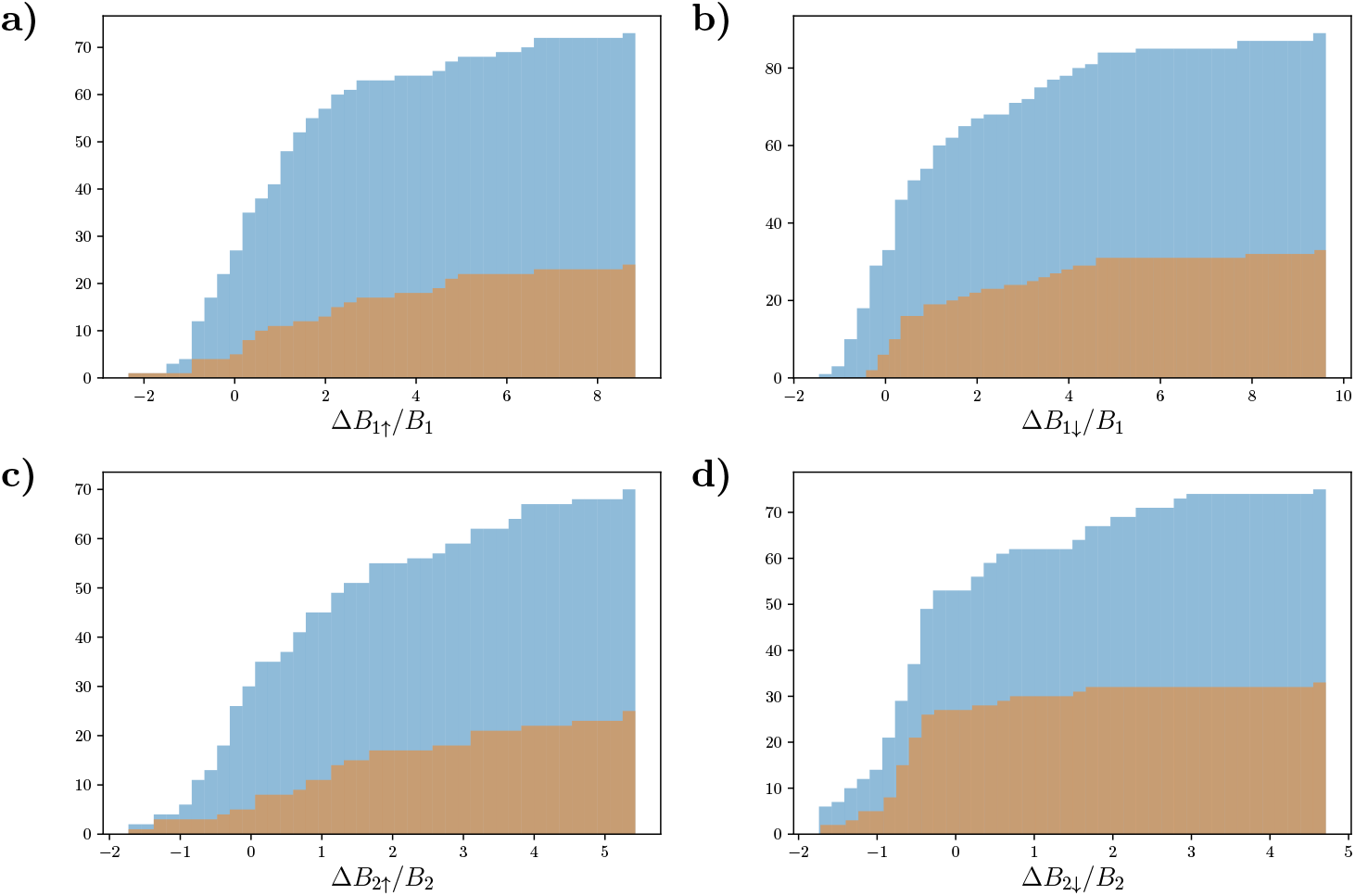
Cumulative distributions of the estimated responses to nutrient enrichment (↑) and fish addition (↓) treatments of each trophic level biomass. Same information as in Figure S1.1 using cumulative distributions. Blue distributions represent the estimated responses of all possible chains analysed before filtering. Orange distributions show the estimated responses of the chains that were identified as dynamically consistent chains. As mentioned above, in dynamically consistent chains, we expect the two levels to respond in the same direction to nutrient enrichment, and in opposite directions to fish addition. Therefore, in (a, b and c) most of the chains selected present a positive biomass response to the perturbation and are accumulated after 0. In contrast, biomass responses of trophic level 2 to fish addition (d) are expected to be generally negative. Thus, we observe that very few chains are selected when trophic level 2 presents a positive response to fish addition, i.e., there is a very small accumulation of chains after 0.

**Figure S2.3:**
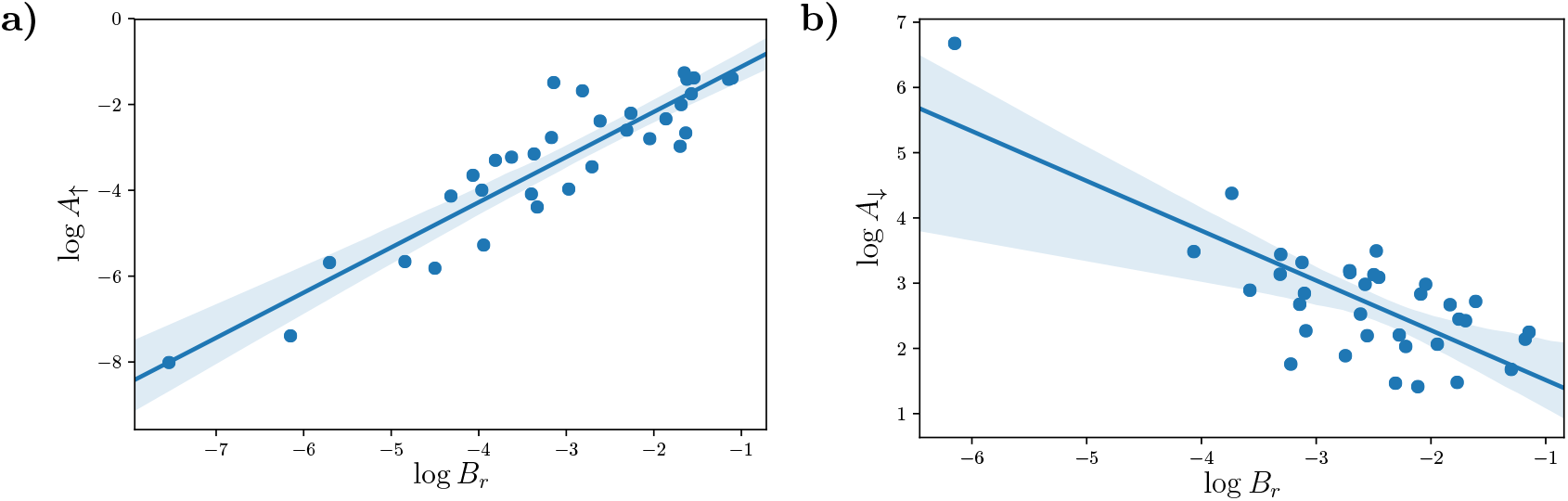
Empirical relationship between biomass distribution across trophic levels and response ratio without exclusion of the outliers. (a) Relationship between biomass ratio 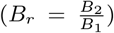 and bottom-up response ratio 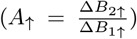 due to nutrient enrichment. (b) Relationship between biomass ratio and top-down response ratio 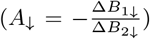 due to fish addition. Each point represents a food chain in our analyses. Blue lines represent the linear regression and shaded areas represent 95% confidence intervals.

### Appendix S3. Supplementary ABC simulation results

**Figure S3.1:**
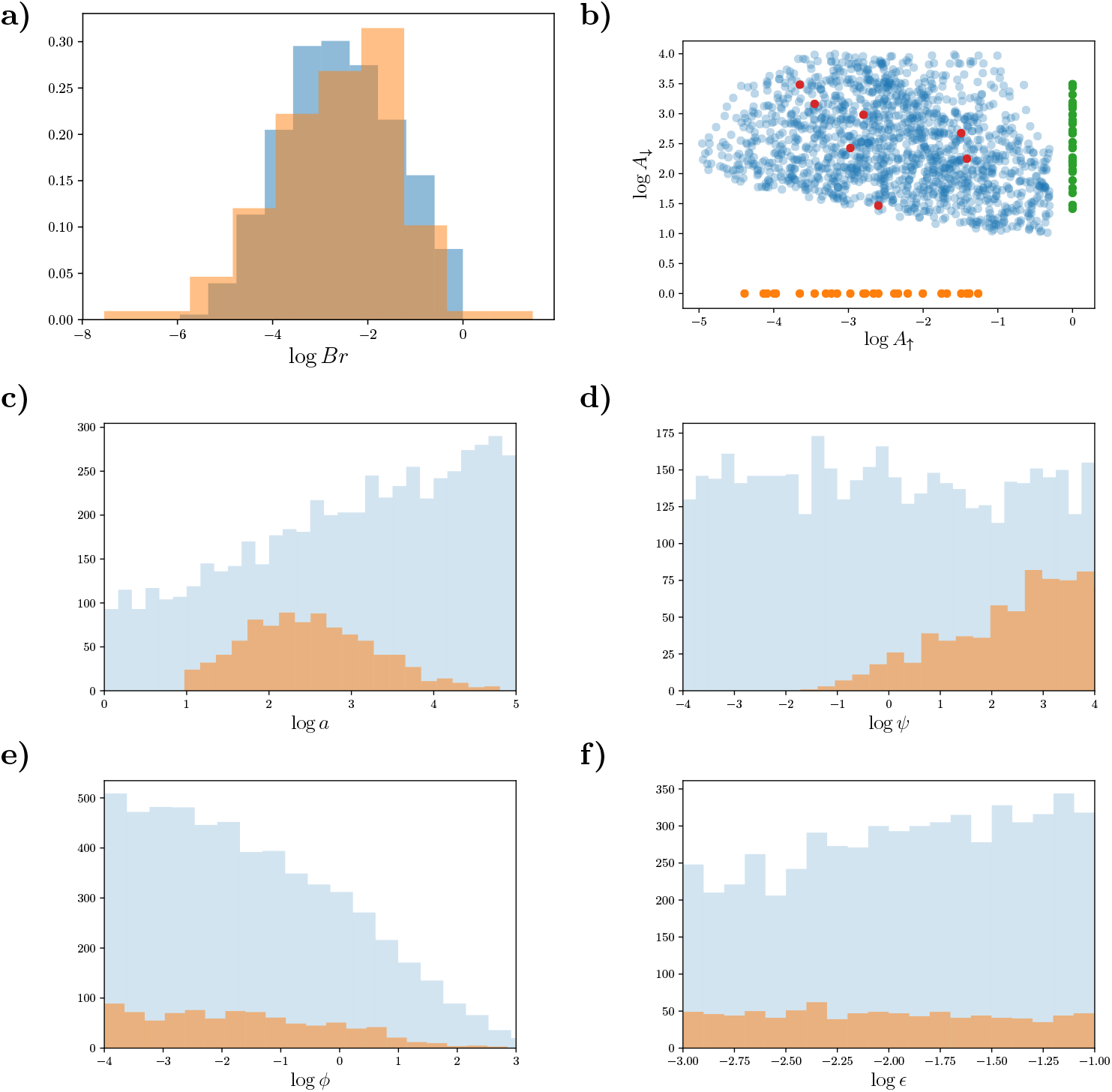
Experimental data versus simulated data. (a) Histograms of the biomass ratio in the experimental systems (orange) and in the simulated data (blue). Both distributions are normalised such that the area under the histogram sums to 1. (b) Relationship between bottom-up amplification 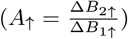 due to nutrient enrichment and top-down amplification 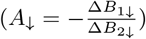 due to fish addition. Orange points represent all the *A*_↑_ values in the empirical data. Green points represent all the *A*_↓_ values in the empirical data. Red points represent the empirical data for which both values of *A*_↑_ and *A*_↓_ were accessible. Blue points represent the simulated data, constrained to reproduce empirical patterns (ABC inference method). Note the slight negative correlation between *A*_↑_ and *A*_↓_. (c) Distribution of relative attack rates *a* = *α/D*_1_ after constraining for coexistence (light blue) and after constraining by empirical patterns (orange). (d) as in c) but for ratio of consumer to producer self-regulation *ψ* = *D*_2_/*D*_1_. The large variation of *ψ* may be responsible for the anticorrelation of *A*_↑_ and *A*_↓_ observed in (b). Indeed *A*_↑_/*A*_↓_ ~ *ψ*. The broadest possible range of *ψ* is achieved if the larger *A*_↑_ are associated with the smallest *A*_↓_ and vice versa. Conversely, the larger the range in *ψ* the stronger the anti-correlation between the two response ratios. (e) as in (c-d) but for consumer losses over primary productivity *ϕ* = *r/g*. (f) as in (c-e) but for conversion efficiency *ε*.

**Figure S3.2:**
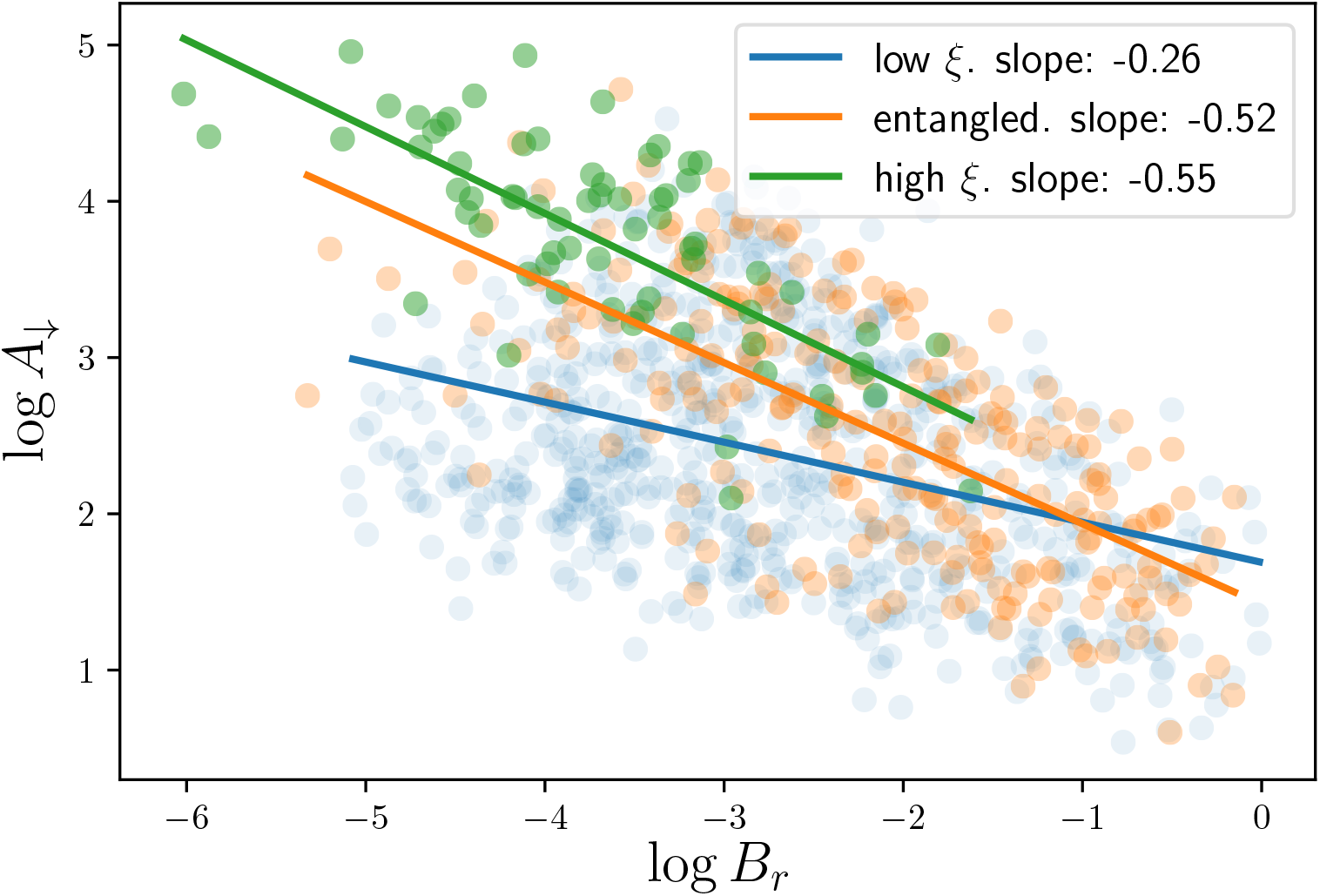
Relationship for *A*_↓_ and *B_r_* for the three regimes of dissipative release, where the qualitatively relationship is maintained but their interpretation should differ. From a correlation for low dissipation release to an actual causation for high dissipation release.

## Notes

### Competing Interest Statement

The authors have declared no competing interest.

### Summary of Updates

Revised manuscript, main results and ideas remain, but the presentation and some interpretations have been improved.

